# Ultrasensitive detection of TCR hypervariable region in solid-tissue RNA-seq data

**DOI:** 10.1101/073395

**Authors:** Bo Li, Taiwen Li, Binbin Wang, Ruoxu Dou, Jean-Chrisophe Pignon, Toni K. Choueiri, Sabina Signoretti, Jun S Liu, Xiaole Shirley Liu

**Affiliations:** Department of Biostatistics and Computational Biology, Dana Farber Cancer Institute; Department of Statistics, Harvard University; State Key Laboratory of Oral Diseases, West China Hospital of Stomatology, Sichuan University; School of Life Science and Technology, Tongji University, China; Department of Colorectal Surgery, Sixth Affiliated Hospital, Sun Yat-sen University; Department of Pathology, Brigham and Women’s Hospital; Kidney Cancer Center, Dana Farber Cancer Institute

## Abstract

Characterization of tissue-infiltrating T cell repertoire is critical to understanding tumor-immune interactions and autoimmune disease etiology. We present TRUST, an open source algorithm for calling the TCR transcript hypervariable CDR3 regions using unselected RNA-seq data profiled from solid tissues. TRUST achieved high sensitivity in CDR3 calling even for samples with low sequencing depth and has demonstrated utilities in its application to large tumor cohorts.

---

Neoantigen-specific tumor-infiltrating T lymphocytes (tTIL) are functional carriers for cancer cell elimination and are the primary focus of immunotherapies^1‐4^. Characterization of the tTIL receptor repertoire is critical to understanding tumor-immune interactions. Profiling of the tTIL β chain hypervariable complementarity-determining region 3 (CDR3) relies on targeted PCR capture followed by deep next generation sequencing (TCR-seq) ^5^ and has recently demonstrated significant clinical impact^6, 7^. However, TCR-seq is unavailable for clinical studies with limited tissue materials, as high-throughput profiling of the tumor genome and transcriptome is usually of higher priority. With the increasing interest to study both tumor and its infiltrating immune repertoire, there is a pressing need to profile tTIL through repurposing tumor RNA-seq data, instead of consuming additional tissues for TCR-seq. Motivated by such need, we developed a computational tool, <u>T</u>CR <u>R</u>epertoire <u>U</u>tilities for <u>S</u>olid <u>T</u>issue, or TRUST, for ultrasensitive detection and de novo assembly of the hypervariable CDR3 regions using RNA-seq data (Online Methods). In this work we show that TRUST achieved recall rate an order of magnitude higher than contemporary methods ^8–10^, even at extremely low coverage of the TCR transcripts.

Given RNA-seq reads previously aligned to the reference genome, TRUST first detects library type (Fig. 1a) and sorts the reads for CDR3 detection (Fig. 1b). In paired-end library (PE) mode, TRUST searches for pairs where one mate is properly mapped to the TCR gene loci and the other mate is unmappable. In single-end library (SE) mode, TRUST directly selects unmapped reads with consensus CDR3 motifs (Fig. 1c). Read pairs with both mates unmappable are subject to the same analysis as single-end reads if they contain CDR3 sequences. Users can also force TRUST to run in single-end mode for paired-end library. Sorted unmapped reads are candidates for CDR3 assembly and are assigned to TCR genes based on their mapped mates (PE) or the partial sequences they contain (SE) (Fig. 1d). For each TCR gene, TRUST assembles reads into contigs based on partial sequence overlap (Fig. 1e). It then realigns the flanking regions of each contig to the IMGT reference sequences ^11^ and reports ones with at least 6 consecutive matched amino acids(Fig. 1f). Finally, TRUST reports the complete or partial CDR3 assemblies and related variable (V) or joining (J) genes in FASTA format.

**Figure 1.**
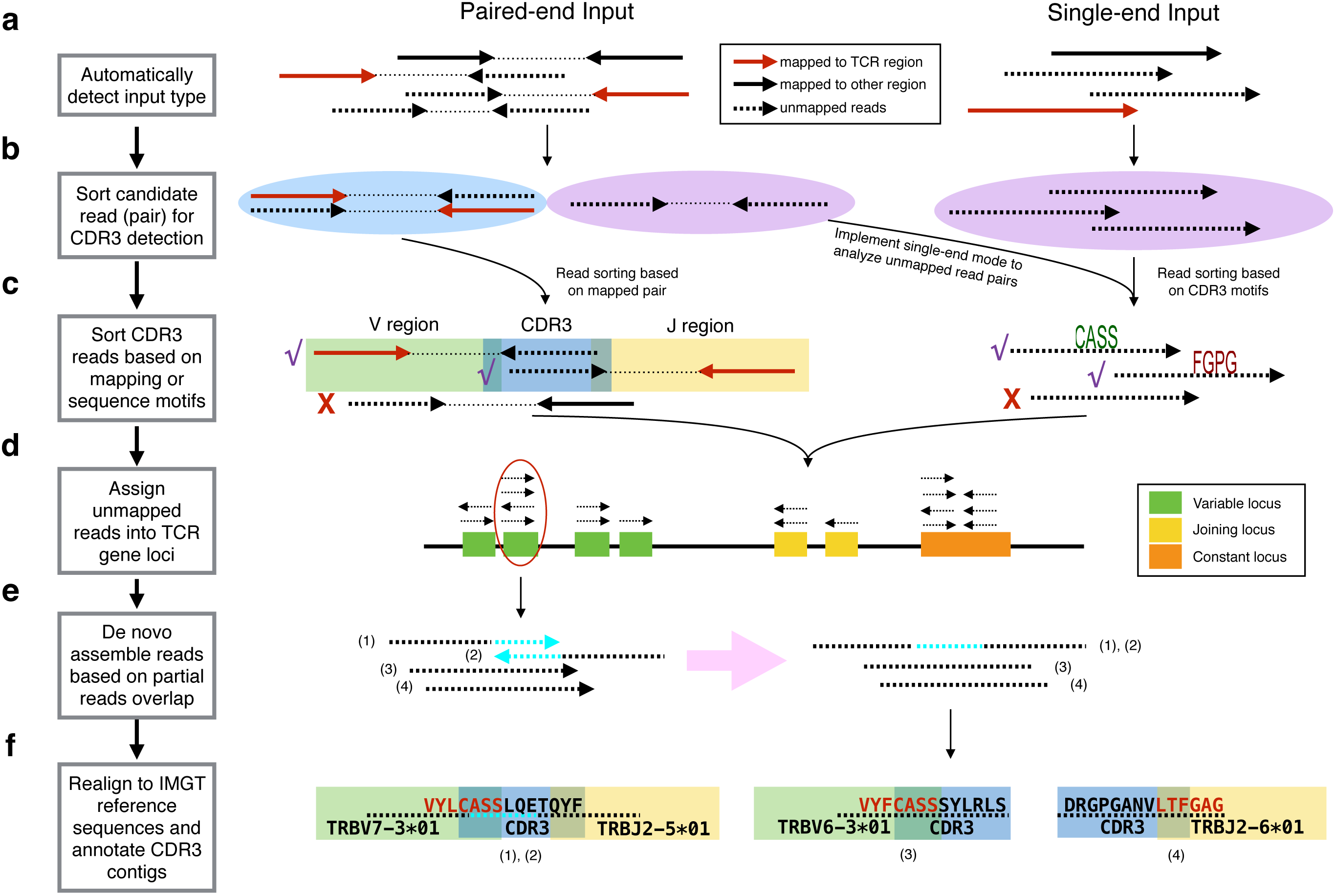
Cartoon summary of TRUST method workflow. TRUST takes the input in BAM format, with reads previous aligned to human reference genome. It automatically determines if input library is paired-end (PE) or single-end (SE) (**a**). For PE input, TRUST selects read pairs with one mate properly mapped (solid arrow) and the other unmapped (dashed arrow). For SE input, TRUST selects all the unmapped reads for further processing (**b**). In the next step, TRUST sort CDR3-related unmapped reads based either of the two criteria: its paired mate properly aligning to the TCR region (red arrow) or the read containing putative CDR3 motifs (**c**). For PE library, each read is assigned to individual TCR genes based on the mapping location of its paired mate. In case of SE library, the read is aligned to the motif-containing regions of the IMGT reference DNA sequences. If the top hit has a maximum mapping score larger than 20, the read is kept and the corresponding TCR gene is assigned (**d**). De novo assembly is then performed for reads assigned to each of the TCR genes individually (**e**). If two reads share no less than N overlapping bases, with up to E mismatch tolerated, they will be joined into one contig. N and E are user-defined parameters, with N default 20 and E default 1. Contigs assembled from the previous step are annotated by re-alignment to the IMGT reference gene amino acid sequences of the motif-flanking region. Red letters in e are mapped template amino acids. Contigs not mapped to any reference gene are discarded. Kept contigs are annotated for the CDR3 sequences and corresponding variable and/or joining genes. TRUST reports variable gene for the N-terminus partial assemblies, joining gene for the C-terminus partial assemblies and both genes to the complete assemblies.

In order to test if TRUST assembles real CDR3 sequences, we performed TCR-seq on the β chain of 3 formalin-fixed paraffin-embedded (FFPE) samples of TCGA kidney clear cell cancer with RNA-seq data available ^12^ (Supplementary Table 1). CDR3s called using TCR-seq are considered as real transcripts of the infiltrating T cells. We ran TRUST in both single-end and paired-end mode on the RNA-seq data of these samples and compared TRUST results with TCR-seq calls (Online Methods). In both modes, approximately two thirds of TRUST assembled CDR3s were found in the TCR-seq data (Fig. 2a and Supplementary Fig. 1a), confirming that TRUST is capable of calling real CDR3 sequences. DNA extracted from FFPE samples often has degradation ^13^ and may not capture the entire tTIL repertoire ^14^. In addition, tissue slides used for RNA-seq and TCR-seq were in different locations and may not contain the same tTIL. Therefore, it is expected that a fraction of TRUST calls were not found in the TCR-seq results. CDR3s called from both TCR-seq and TRUST are enriched for high frequency clonotypes (Fig. 2b and Supplementary Figure 1b). Specifically, TRUST identified an average of 53.5% of the top 1% most abundant CDR3s from TCR-seq. In addition, V/J gene assignments by TRUST are highly concordant with those obtained from TCR-seq (Fig. 2c and Supplementary Figure 1c).

**Figure 2.**
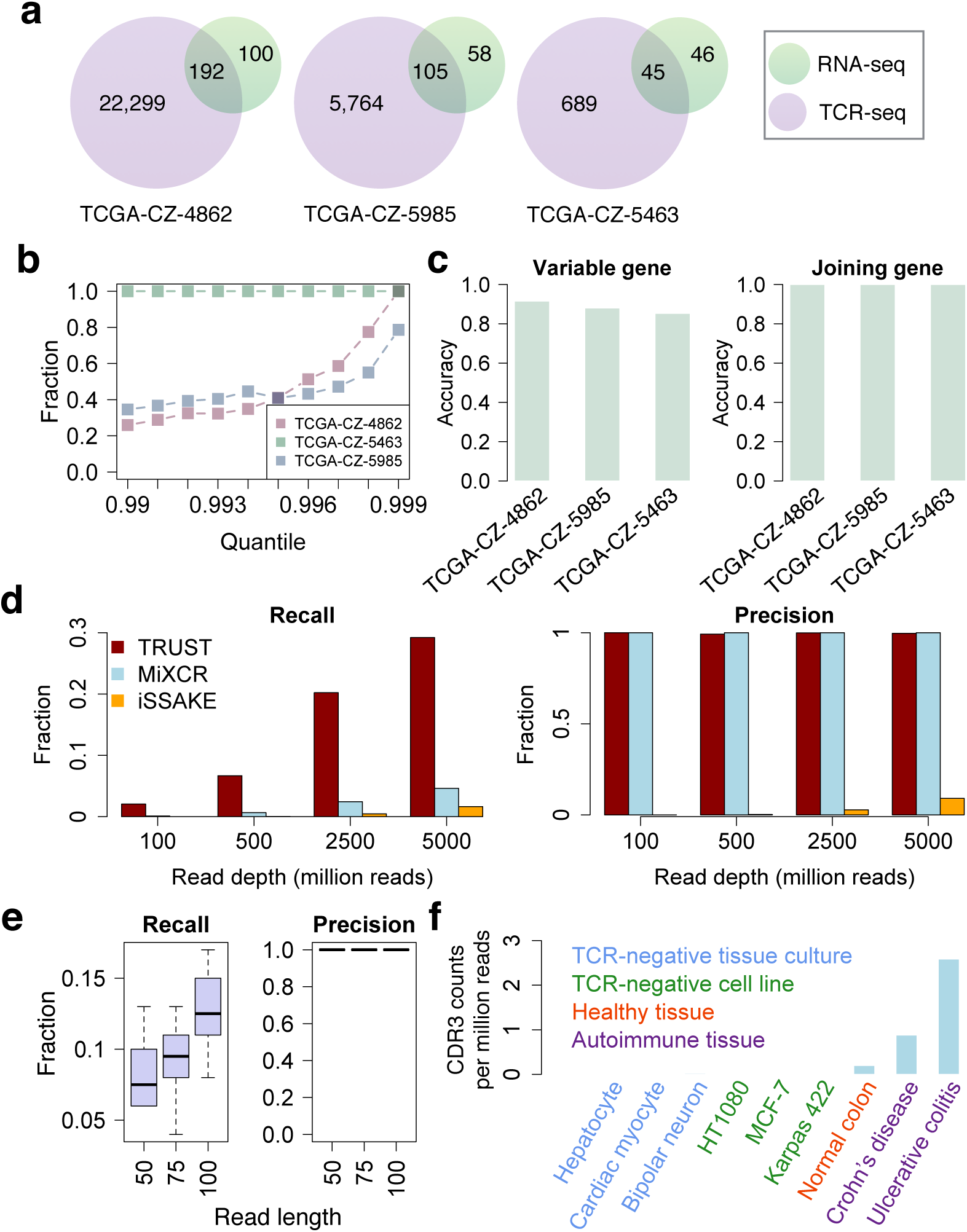
Performance evaluation of TRUST in single-end mode. **a**. Venn diagrams showing the number of CDR3 sequences called from TCR-seq (ImmunoSeq), RNA-seq using TRUST and the overlap of the two approaches for 3 TCGA kidney renal clear cell carcinoma samples. **b**. TRUST-reported CDR3s are enriched for clonotypes with high abundance. At each quantile (Q), Y-axis is the fraction of TRUST-reported CDR3 with clonal frequency above or equal to Q. Same analysis was repeated for all three TCGA samples. **c**. Accuracy of variable/joining gene estimations by TRUST. Accuracy is the percentage of correctedly assigned variable or joining genes by TRUST using the overlapping CDR3s (a) and TCR-seq reports as true positive. **d**. Recall and precision estimations based on *in silico* simulations at different read depths. 100 TCRβ transcripts were randomly sampled using IMGT reference genes, mimicking the process of V(D)J recombination. 100 simulations were performed for each of the four different coverage settings, 0.02, 0.1, 0.5, 1.0, which correspond to read depths of 100M, 500M, 2,500M and 5,000M respectively. TRUST, MiXCR and iSSAKE were applied to the simulated datasets. Recall and precision were estimated by comparing to the original 100 sampled TCRβ transcripts. **e**. Recall and precision estimations at different read length settings. Similar simulations to **d** were performed for libraries size 500M with 50, 75 and 100nt reads. Recalls are displayed as lower boxes and precisions are upper boxes. **f**. Application of TRUST to non-cancerous tissue samples. Nine single-end RNA-seq samples from the public domain were collected. These samples fall into four categories, based on their TCR-containing and disease status. Y-axis is the number of TRUST-reported CDR3 normalized by the read depth of input data.

Since TCR-seq from FFPE samples might have limited coverage, we implemented *in silico* simulations to thoroughly evaluate the performance of TRUST. To this end, we generated short read sequencing data that resembles real RNA-seq data, with known true positive TCR transcripts (Supplementary Figure 2 and Online Methods). It is estimated that in an RNA-seq data with 200 million (M) 50nt reads, the coverage for TCR transcripts is 0.04 ^12^. We first simulated SE library with read length 50nt and read depth from 100M to 5,000M, which is equivalent to coverage from 0.02 to 1.0. Read depth refers to the total number of reads in an RNA-seq data. For PE library, read depth is twice as the number of DNA fragment, which contains two reads produced from a same haplotype. We randomly sampled 100 TCRβ transcripts for each simulation and performed 100 simulations at each coverage. TRUST achieved average recall of 2.1% at read depth 100M, while two competing methods MiXCR ^10^ and iSSAKE ^8^ were 0.12% and 0.0% respectively at the same read depth. Precisions of TRUST, MiXCR and iSSAKE at read depth 100M were 100%, 100% and 0.0% (Fig. 2d). Similar recall and precision were observed for PE library simulations as well (Supplementary Figure 3). We also applied Decombinator ^9^ on the simulated data, but it failed to assemble any contig even at read depth 5,000M. The above results suggest that TRUST is a superior method suitable for calling CDR3s from RNA-seq data with better sensitivity and specificity than other contemporary methods.

We performed another set of simulations to investigate TRUST performance over different read lengths. Fixing read depth at 500M, we simulated libraries with read length 50, 75 and 100nt (Online Methods). The TRUST recall increases with longer reads, with precision not influenced by either SE or PE inputs (Fig. 2e and Supplementary Figure 4). This is because long reads have higher possibility to cover the CDR3 region than shorter reads, when the number of fragments is fixed. The fraction of complete calls (defined as CDR3 assemblies containing both N- and C-terminus motifs) also increases with longer reads, and this fraction is unaffected by read depth (Supplementary Figure 5).

Non-cancerous solid tissues sometimes also contain a sizeable amount of infiltrating T cells, especially in case of autoimmune diseases. In order to explore the utilities of TRUST on other types of solid tissues, we collected 9 samples from the public domain, including 6 negative control cell line samples from ENCODE ^15^ and 3 solid tissues from NCBI sequence read archive (SRA). Of the 6 cell line samples, 3 of them are in vivo differentiated normal cells and 3 are cancer cell lines. Since V(D)J recombination only occurs in the lymphocyte lineage, these 6 cell lines are expected to be TCR-negative. Using CDR3 counts per million reads (CPM) as a measure of TCR content, we found CPMs to be extremely low or zero in the cell lines as expected. We then compared the TCR contents in the normal colon and colon tissues with Crohn’s disease or ulcerative colitis, and found the amount of T cells to be much higher in inflamed tissues than normal tissue (Fig. 2f and Supplementary Table 2). As TRUST assembles an appreciable amount of CDR3 sequences even at very low sequencing depth, these results suggest that TRUST is potentially suitable for analyzing RNA-seq data profiled from non-cancerous inflammatory solid tissues as well.

A prototype of TRUST has been applied to a large cohort of cancer samples and revealed rich information of tumor-immune interactions ^12^. The current version of TRUST is built on this prototype with three important improvements. First, while the prototype only handles paired-end data, TRUST also analyzes single-end library RNA-seq data with comparable performance as paired-end input. Second, the inclusion of unmapped read pairs (Fig. 1b) as well as optimized CDR3 realignment and annotation approach increased the recall by 91% (4.2% at 100M read depth, PE mode compared to the prototype 2.2%) while maintaining high precision. Third, the coverage for high abundance T cell clones (top 1% quantile) is increased from 12.8% to 58.5% (PE mode). It is known that antigen-specific infiltrating T cells undergo clonal expansion and become more abundant in the repertoire ^16^. Further more, recent study revealed that high T cell clonality is associated with improved outcome in anti-PD1 therapies in melanoma ^6^. Therefore, the high abundance T cells recovered by TRUST may harbor tumor-reactive clones with direct clinical applications.

In summary, we developed TRUST for ultrasensitive calling of CDR3 sequences using unselected RNA-seq data profiled from solid tissues which is applicable to both single-end and paired-end sequencing libraries. TRUST has demonstrated utilities in application to large cancer sample cohorts and is a unique addition to the current sequence analysis toolset. In this work, we observed an average recall of 2-4% for RNA-seq data with 100M reads. Recent technology development has enabled the sequencing of PE 150nt library of 300M DNA fragments (600M reads) inexpensively ^17^. At this setting, TRUST can achieve 9.4% average recall with 96.6% precision and 76.6% complete CDR3 assembly rate (Supplementary Figure 6). Therefore, approximately one tenth of the TCR repertoire can be accurately retrieved by repurposing the RNA-seq data. With rapidly accumulating RNA-seq data and continuously decreasing sequencing cost, we anticipate TRUST to attract broad interest in the immunology and cancer research fields.

## Methods

### TCR hypervariable region de novo assembly workflow

#### i). Automatic detection of input library type

TRUST implements different approaches of read sorting and alignment for singled-end and paired-end libraries and therefore, the first step in the method is to detect library type. Given the input BAM file, TRUST initiates a read screen and selects unmapped reads. If the no paired-end fragments (first bit of SAM flag is 0) in the first million reads, TRUST assigns single-end library to the input data. Oppositely, if paired-end fragments are detected in the first million reads, TRUST assigns paired-end library to the input data. User can force TRUST to run in single-end mode (-s option) for paired-end library data. When this option is enabled, TRUST ignores paired-end information and always treats input data as single-end.

#### ii). Read sorting for CDR3 detection

After input library type is determined, TRUST searches the entire library for informative read or read pairs. For paired-end library, TRUST keeps read pairs with at least one mate unmapped to the reference genome. For single-end library, TRUST keeps the unmapped reads. A subset of these informative read or read pairs may contain the hypervariable CDR3 region and cannot map to the reference genome due to insertion of non-template DNA bases.

#### iii). Sorting reads based on mapping or consensus motifs

For paired-end library input, TRUST finds unmapped read with its paired mate properly mapped to the TCR gene regions, including variable, joining and constant genes. The genomics locations for the TCR loci are (hg19 assembly): α (chr14:22090057-23021075), β (chr7:141998851-142510972), γ (chr7:38279625-38407656). δ gene region is embedded in the α region (chr14:22891537-22935569) and therefore the reads will be obtained along with TCRα. For single-end library input, TRUST determines if a read is potentially informative by whether it contains CDR3 DNA consensus motifs. The motifs are predefined based on IMGT variable or joining gene sequences. For paired-end library, if both mates are unmapped, they could potentially contain CDR3 region as well. These reads are subject to TRUST single-end library analysis. Redundant contigs resulted from paired reads of same fragment are removed in the downstream analysis.

#### iv). Sort CDR3 reads into TCR genes

As TRUST uses a comprehensive pairwise read comparison method for sequence assembly, its time cost is O(N^2^), where N is the number of candidate CDR3 reads. We here use a divide and conquer approach to reduce the time cost by assigning CDR3 reads into smaller sets and process each set individually. Intuitively, we assign candidate CDR3 reads selected from the previous step into TCR genes. Reads assigned to different genes belonging to the same category (variable or joining) must not come from a same transcript. Therefore, these reads can be assembled individual to TCR genes without losing CDR3 detection power. For paired-end genes, TRUST assigns TCR genes based on their mapped mates. For single-end reads, TRUST aligns the read depending on the type of motif it contains. For example, reads containing β variable gene motifs will be aligned to IMGT β variable gene DNA sequences. If the mapping score of the top hit is greater than 20, the read is kept and assigned to the corresponding gene.

#### v). De novo read assembly based on partial sequence overlap

For unmapped reads assigned to a TCR gene, TRUST performs pairwise sequence comparisons, searching for reads with more than N bases overlap, where N is a user-defined threshold with a default of 10. TRUST uses a two-bit coding for the DNA nucleotide sequences to accelerate computation and reduce memory use. TRUST allows for up to E mismatches in the overlap sequence to tolerate sequencing errors (E is user-defined parameter with a default of 1). For each TCR gene, a read-sharing matrix is produced documenting the sequence overlap information of all reads assigned to this gene.

We use an undirected graph to represent the read-sharing matrix, with each node representing an unmapped read and each edge indicating partial sequence overlap. TRUST then traverses the graph to find all the disjoint subgraphs, or cliques and order reads within each clique based on their overlapping sequences. For example, when the 3’ end of read X overlaps with the 5’ end of read Y, TRUST places X in the 5’ end of Y, and vice versa. If there are reverse complement reads, all the reads are converted to one randomly chosen direction before ordering. After ordering, reads are assembled into one contig. When two reads positioned after a same read do not overlap, they may come from two contigs sharing the same upstream variable gene. In this situation, TRUST assembles each CDR3 contig individually.

As above mentioned, reads assigned to different genes of the same category (variable or joining) must be generated from different TCR transcripts. However it is possible that contigs assigned to different genes of different categories were parts of the same transcript. For example, a contig assembled from a variable gene may be the same transcript as a contig assembled from a joining gene. We define variable gene as ‘upstream genes’ and joining or constant genes as ‘downstream genes’. TRUST then compares each of the upstream/downstream contig pairs, and merge them if they overlap by at least 2N bases, with up to E mismatch(es). Meanwhile, if two contigs are products of the same read pair, one of them are removed to reduce redundancy.

#### vi). Realign contigs to IMGT reference genes and annotate CDR3 regions

The final contigs assembled from above steps must contain at least one CDR3 consensus motif(s) and are likely CDR3 assemblies. For each contig, TRUST first convert the DNA sequence into amino acid (AA) sequence and locates the consensus AA CDR3 motif using a predefined list of patterns. The N terminus residual sequence of a variable motif, or the C terminus residual sequence of a joining motif, is called a flanking region. TRUST realigns the flanking region to the IMGT reference gene amino acid sequences. If the top hit contains at least 6 consecutive matched AAs, including the CDR3 motif, TRUST reports this contig as a CDR3 assembly, and reports its corresponding variable and/or joining gene with highest alignment score. If not, this contig is discarded to reduce false positive rate. The final output includes the input file name, CDR3 amino acid sequence, CDR3 DNA sequence and complete DNA sequence of the assembled contig, in FASTA format.

### In silico simulation

We performed *in silico* simulations to systematically evaluate the performance of TRUST (Supplementary Figure 2). We randomly selected the DNA sequences of 3 β variable (11-1*01, 123*01 and 2*03), 3 β joining (1-1*01, 2-1*01 and 2-4*01) and β constant gene (TRBC1) from IMGT reference sequences and performed *in silico* VJ recombination to generate full-length TCRβ transcripts. Non-template random DNA bases were inserted into the VJ junction, with number of bases follow Poisson(5). Meanwhile, N bases at the 3’end of the variable gene and 5’end of the joining gene were deleted, with N following Poisson(1). 100 transcripts were generated as the true TCR transcripts.

We applied simNGS, a toolbox for simulating next-generation sequencing libraries to produce the *in silico* datasets^18^ using the 100 simulated TCR transcripts. For SE libraries, we directly sampled DNA fragments with given read length (r). For PE libraries, we first sampled DNA fragments with median insert size 2×r+50. Illumina short reads with length r were subsequently sampled from the DNA fragment library. Tophat (v2.1)^19^ was applied to map the short RNA-seq reads (Bowtie2 version 2.2.1)^20^ to the human reference genome (hg19), with default parameters except for disabling coverage search (-no-coverage-search) to accelerate mapping. SAMtools^21^ was applied to generate the corresponding BAM index files. TRUST was then applied to the BAM files for CDR3 detection. CDR3 sequences reported by TRUST were compared to the original 100 simulated TCR transcripts for recall and precision estimations. For coverage evaluation, we set the simLibrary −x option to be 0.02, 0.1, 0.5 and 1.0 for read depth 100M, 500M, 2,500M and 5,000M. 100 simulations were performed for each coverage setting and we fixed read length to be 50nt. For read length evaluation, we set the simLibrary −r option to be 50, 75, 100 and 150nt. 10 simulations were performed for each read length setting. For read length 50, 75 and 100nt, we fixed the number of fragments to be 4 (simLibrary −n 4) for SE library and 2 for PE library, which is equivalent to read depth 500M. For read length 150nt PE library simulations, we fixed coverage 0.36 (simLibrary −x 0.36), which is equivalent to read depth 300M.

### Application of MiXCR, iSSAKE and Decombinator to simulated datasets

We applied default settings of MiXCR, iSSAKE and Decombinator to the *in silico* RNA-seq datasets generated as above described. Outcomes of all 3 methods were compared to the simulated TCR transcripts for performance evaluation. No transcripts were reported at read length 50nt by Decombinator even at read depth 5,000M and therefore it excluded in Fig. 2d.

### Access and analysis of public datasets

RNA sequencing data in fastq format for 6 cell line samples (ENCSR859HWB, ENCSR294NDO, ENCSR023VVO, ENCFF002DHV, ENCFF002DKJ and ENCFF002DLL) were obtained from ENCODE data portal (https://www.encodeproject.org). Other 3 samples of colon normal or diseased tissues (SRR1813898, SRR1813883 and SRR2314045) were downloaded from the NCBI Short Read Archive (SRA) database (http://www.ncbi.nlm.nih.gov/sra) in fastq format. All 9 samples were PE RNA-seq libraries. In order to evaluate the single-end mode performance, we only used half of the original data. All files were first aligned to the human reference genome hg19 using Tophat^19^ with default settings. BAM index files were built using SAMtools^21^. The resulting files, BAM with index, are desired input for TRUST method.

## Author contributions

BL conceived the project, developed the TRUST method and carried out the computational analysis. TL and RD contributed to applications to non-cancerous biological samples. BW contributed to data collection. JCP, SS and TKC performed experimental validation using TCGA KIRC samples. JSL provided expertise on computational analysis and algorithm design. XSL supervised and study and wrote the manuscript with BL.

## Acknowledgements

We acknowledge the following funding sources for supporting our work:NCI grant 1U01 CA180980 and National Natural Science Foundation of China grant 31329003 (to XSL); the Trust family, Loker Pinard, and Michael Brigham Funds for Kidney Cancer Research (to TKC); NIH/NCI DF/HCC Kidney Cancer SPORE P50 CA101942 (to SS and TKC); National Natural Science Foundations of China 81321002 (to TL).

## Code and Data Availability

TRUST software, supporting data and usage are available along this manuscript as a Supplementary Software.

## Supplementary Figure Legends

**Supplementary Figure 1.**
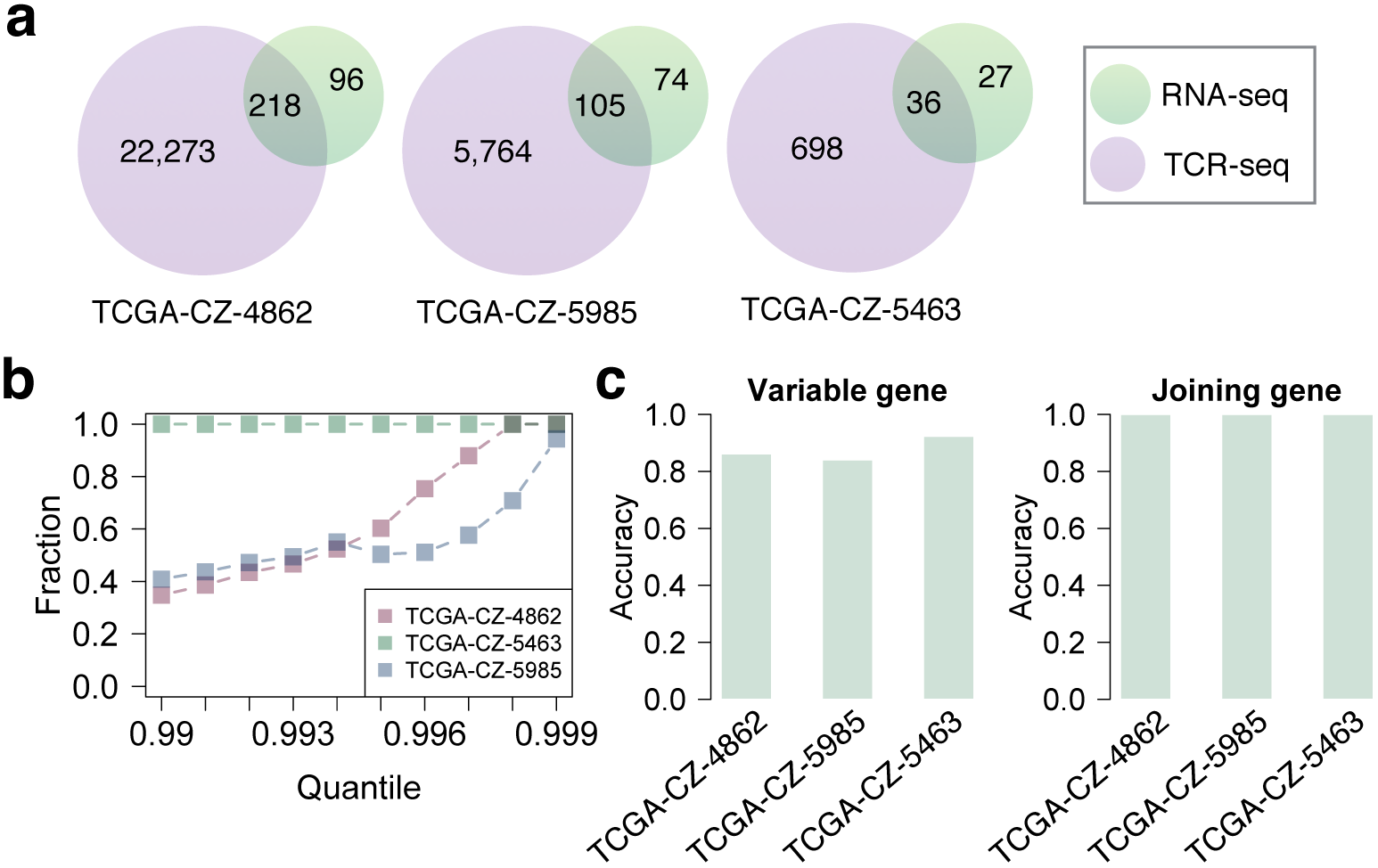
Comparison of TCR-seq results with TRUST-reported CDR3s using RNA-seq data in paired-end mode. Same results are presented as Figure 2a-c, except that TRUST was run in paired-end mode. The performance of TRUST in PE mode is similar to that in SE mode.

**Supplementary Figure 2.**
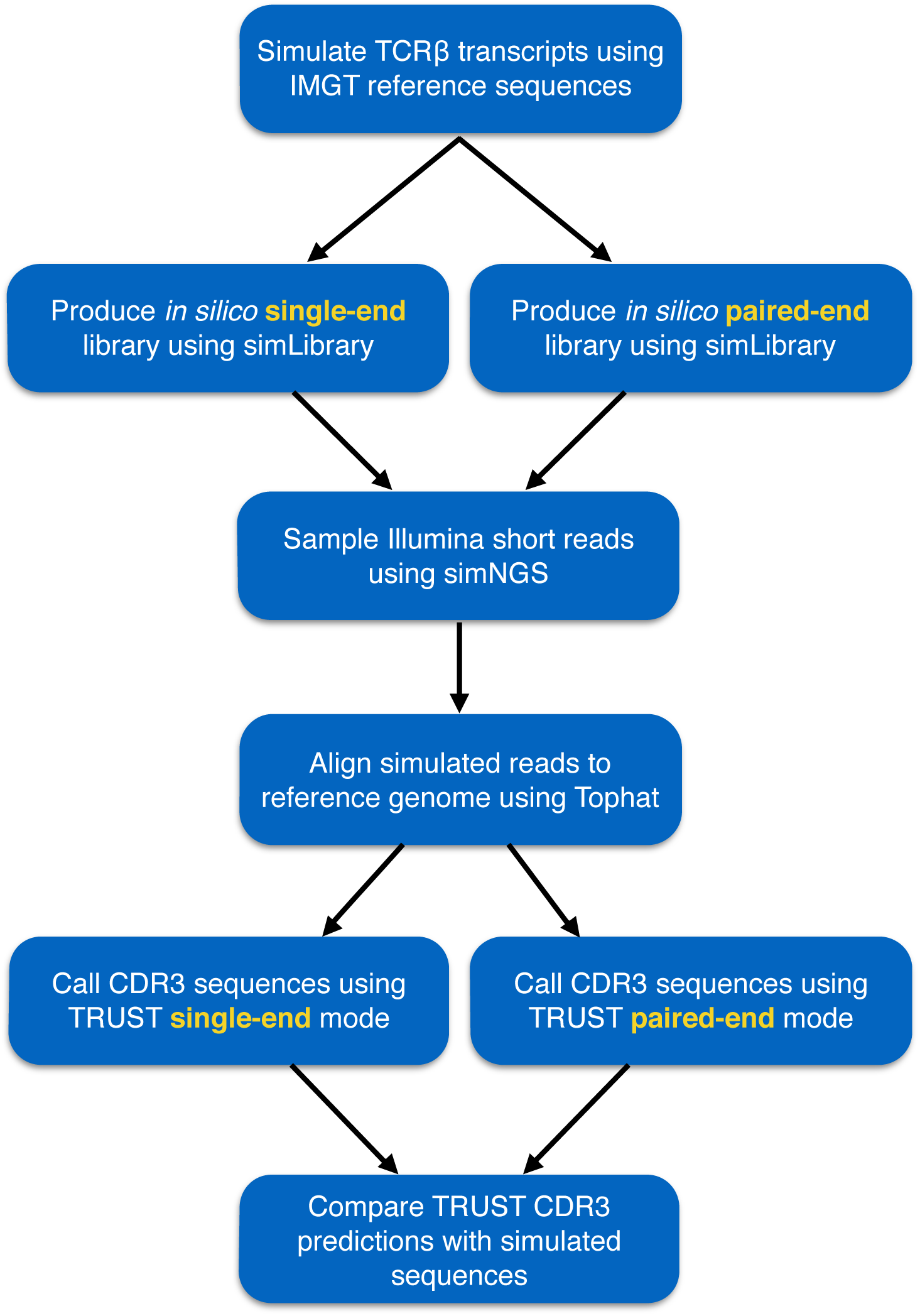
Flowchart of the *in silico* simulation pipeline used for TRUST method evaluation.

**Supplementary Figure 3.**
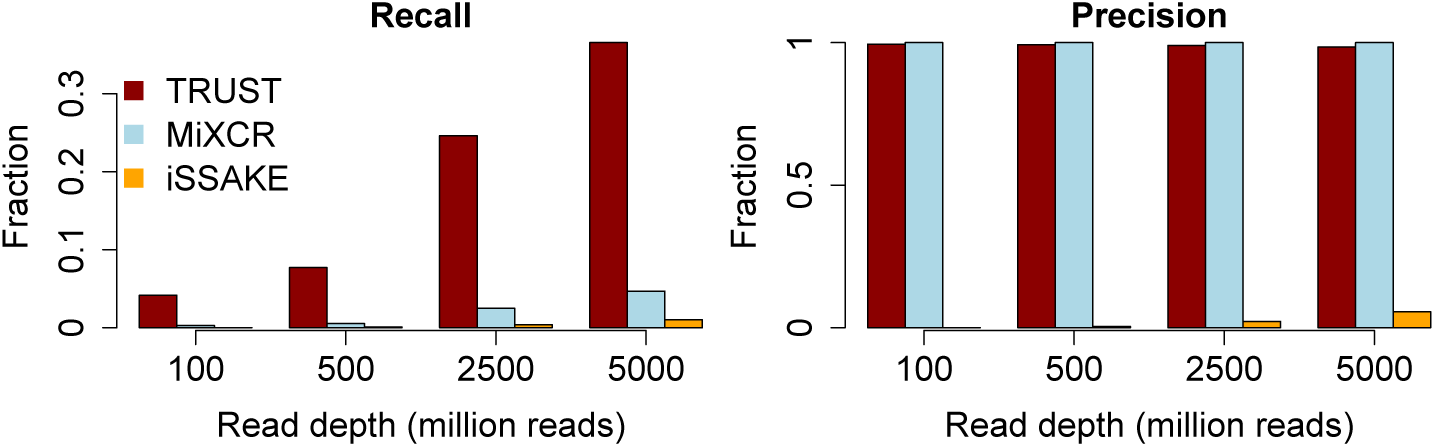
Recall and precision estimations at different read depths in paired-end mode. Similar as Figure 2d, performances of TRUST, MiXCR and iSSAKE were tested on simulated datasets with paired-end libraries. Similar levels of recall and precision were observed for TRUST in PE mode as those in SE mode.

**Supplementary Figure 4.**
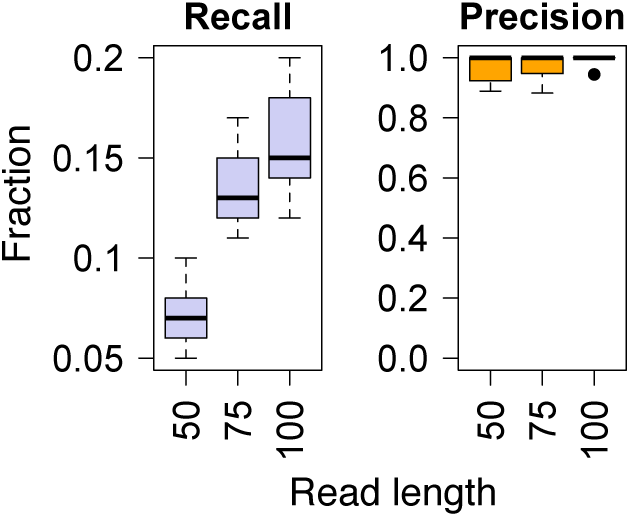
Recall and precision estimations at different read lengths. Paired-end libraries with different read lengths were simulated and the performances of three methods were tested. Similar to single-end libraries, recall and precision were unaffected by read lengths.

**Supplementary Figure 5.**
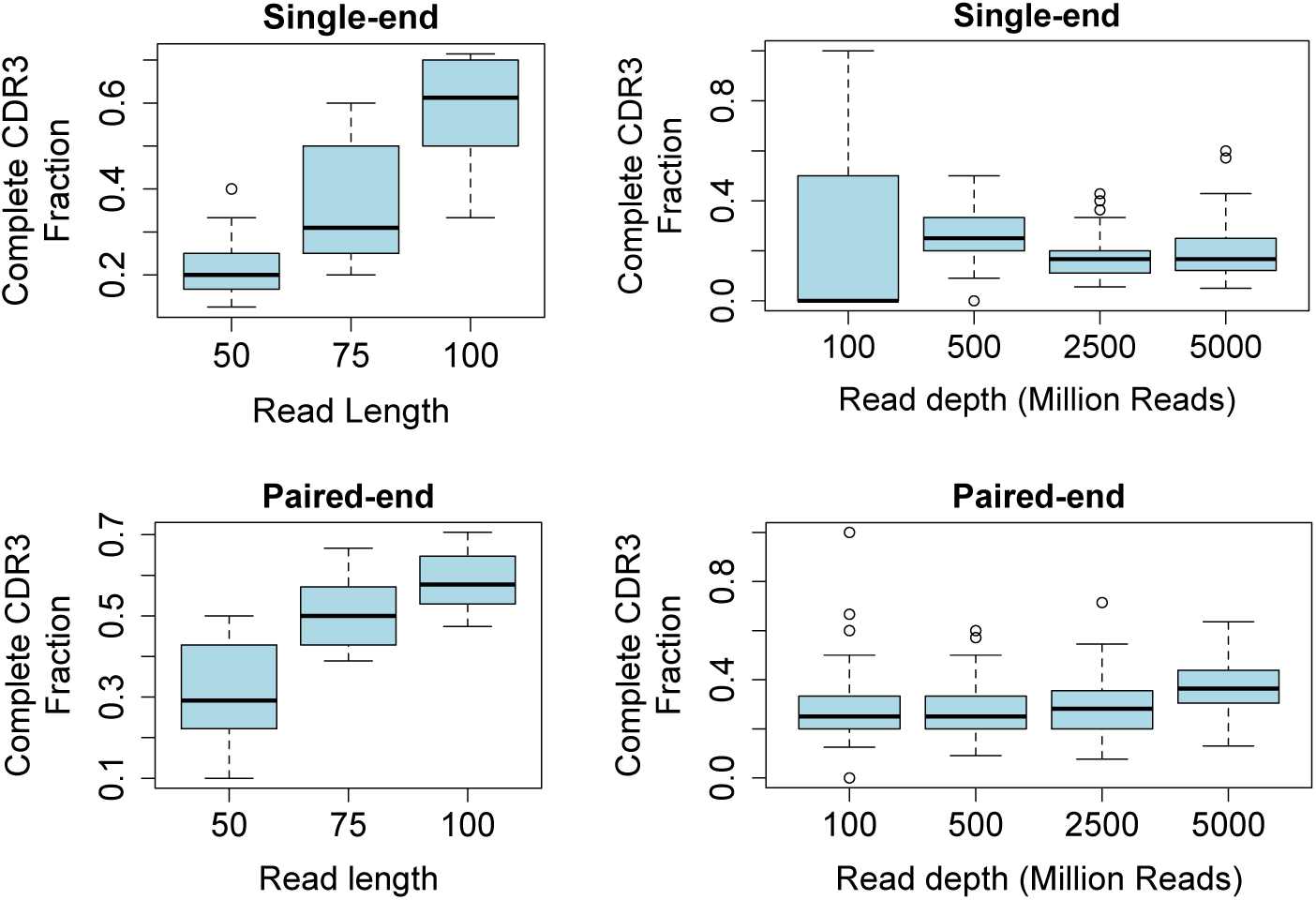
Fraction of complete CDR3 calls with different read length and read depth settings for both single-end and paired-end libraries. Complete CDR3 call rates increase with read length but not read depth for both library types.

**Supplementary Figure 6.**
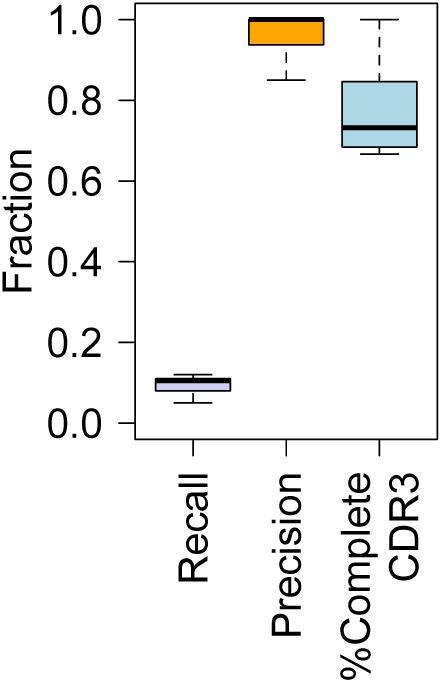
Performance evaluation on an RNA-seq with PE 150nt read length and 300M DNA fragments.

